# Identifying Positional Orthologs by Linear Programming

**DOI:** 10.1101/2025.05.22.655535

**Authors:** Askar Gafurov, Krister M. Swenson

**Affiliations:** CNRS, Montpellier, France

**Keywords:** Comparative Genomics, Positional Orthology, Maximum Disjoint k-Clique, Minimum Orthogonal Partition, Colorful Components, Integer Linear Programming

## Abstract

Studying the evolution of gene orders is essential to our understanding of the link between gene regulation and phenotype. Comparative studies concerning the full complement of genes in large numbers of genomes are now possible, thanks to the advent of affordable long-read sequencing and whole chromosome assembly, combined with automated genome annotation. The study of gene orders remains cumbersome, however, due to a lack of a streamlined, and standardized, process meant for the identification of *positional orthologs*. Within a family of homologous genes, these are those pairs of orthologous gene copies that descend from the same locus in their most recent ancestor, not having been produced by an intervening duplication event. This article approaches the detection of positional orthologs by way of the MAXIMUM COLORFUL GRAPH PARTITION Problem. We present two novel integer linear programming solutions to this problem, evaluate their efficacy on simulated and real data, while comparing them to several new and existing heuristics, as well as to the Double Cut and Join based ILP called `FFGC`.

**Digital Object Identifier:** 10.4230/LIPIcs.CVIT.2016.23

## 1 Introduction

Gene organization on the chromosomes have enormous phenotypic and evolutionary consequences. Genome rearrangements, also known as *structural variations*, play an important role in reproductive isolation [24], and in adaptation [18]. Study of this role in its infancy, yet it is so significant that a recent special issue of Molecular Ecology was recently devoted to the subject [1]. The regulatory effects of rearrangements are most dramatically present in their disease-causing form [10, 17].

Yet genome rearrangements remain extremely difficult to study. With the advent of long-read sequencing and Hi-C, the proper assembly of chromosomes is only now becoming possible on a large scale; the very mechanisms that create a rearrangement are also those that create repetitive regions that are difficult to assemble. These sequences then need to be annotated with gene annotation pipelines, or with whole genome alignment (WGA) methods. While the WGA methods use, by nature, both the nucleotide similarity and the genomic context around a region, the gene annotation methods group genes into homologous sets, called *gene families*. These sets often contain multiple copies of a gene for each gene family. Due to duplicate genes, further processing is required before using these gene families in down-stream analyses. This step often entails partitioning the set into clusters of orthologous genes (COGs). The Quest for Orthologs consortium [14] is dedicated to the development and comparison [3] of these methods. Due to their sequence alignment-based methodologies, these studies have the goal of inferring single-copy orthology sets that have the highest sequence similarity. These sets are most useful for phylogenetic studies, although care must be taken in their proper use [23].

On the other hand, the gene copies that have inherited their position through evolution, are those that are the most important for gene-order studies [8]. These are not the same as COGs, as one of the duplicated genes may rapidly mutate, independent of genomic position. Downstream gene order analyses using orthologous groups inferred from sequence similarity alone will certainly be confused by orthologs sets inferred via typical means.

It is therefore imperative to infer sets of *positional orthologs*, which are sets of orthologs where each pair of genes in the set can be traced to their ancestor without having been the target of a gene duplication along that path. For a clear presentation of the concepts concerning positional orthology, see Dewey [8]. In the case of what Dewey calls *asymmetric* duplication, where the original gene copy can be unambiguously identified directly after the duplication event, the gene neighborhood after the duplication can subsequently be used to infer positional orthology. In the case of *symmetric* duplication such as tandem duplication, where the target is not obvious, one must rely on sequence similarity alone.

Using gene context for orthology inference has its roots in Sankoff’s exemplar problem for gene orders which, given two gene orders, asks for the version of those orders containing only a single copy of a gene from each family, such that the rearrangement distance is minimized [21]. The first large-scale software for the inference of positional orthologs that minimized a genome rearrangement distance was `SOAR,` later extended to multiple genomes in `MultiMSOAR`[22].

Jun et al. [13] introduced a method for identifying positional orthologs between two genomes. It is based on the assignment problem, with gene pairs weighted by sequence similarity, and edges filtered using a threshold for shared genes in adjacent neighborhoods. Two related software were since developed, using a similar setup. `OrthoGNC` added additional neighborhood filters [12], while `BBH-LS` added a bidirectional-best-hits criterion to matching process [25]. Concurrently, `EGM` was introduced with a similar bipartite matching concept coupled with gene-pair weights that are a function of the sequence similarities between the neighboring genes. All of these methods may be considered to be “reference based”, given their pair-wise nature.

`Poff` [16] combines Proteinortho [15] and `FFAdj-MCS` [9], the former being used to cluster genes into families and weight gene pairs, and the latter being used to choose orthologs based on adjacent gene information. More recently `FFGC,` a generalization of `FFAdj-MCS,` has been developed [19, 4]. It assigns positional orthologs with respect to a weighted sum of the double cut and join (DCJ) insertion/deletion rearrangement distance and the similarity scores within and between each positional ortholog set.

In this article, we present a new method for the inference of positional ortholog sets simultaneously in all genomes. The method is based on the Maximum Colorful Graph Partition problem, which has previously been used for orthology prediction based solely on sequence similarity [26]. While being NP-Hard, it admits a 1/*k*-approximation, where *k* is the number of genomes, for an *k*-clique equivalent formulation [11]. We adapt the problem to our purposes by weighting the edges of the graph by genomic neighborhood context in addition to sequence similarity.

We present two novel ILP formulations for MCGP and extensively evaluate the accuracy of our results using the state-of-the-art `Zombi` whole genome simulator [7]. We compare our performance to that of `FFGC,` as it is a generalization of `FFAdj-MCS.` Since it typically takes a set of complete genomes *without* gene family information, the amount computation can be prohibitive. Therefore, we use a modified version of the program that limits its assignments to gene pairs within gene families. We avoid the other methods, many of which were ahead of their time, including `MultiMSOAR, BBH-LS`, and Jun et al., all being over a decade old and either not actively maintained, or unavailable. `OrthoGNC` is very poorly documented, and `EGM` is limited to pair-wise comparisons.

We show that solutions given by MCGP are more accurate than `FFGC,` and that crucially, use of genomic context in the input to the MCGP problem is essential to this accuracy. While the ILP does not always converge in the allotted time, it is almost always more accurate than the approximation algorithm of He et al. [11]. Finally, we apply our methods to four newly assembled *Anopheles* genomes.

### 1.1 Definitions

A chromosome, be it linear or circular, is a sequence of genes, called a *gene order*. Genes that descend from a common evolutionary ancestor are called *homologs*. Homologs that arose from a speciation event are called *orthologs*. Homologs that arose from a duplication event are called *paralogs*. A pair of orthologs are *positional orthologs* when each of them is the descendant of the original gene copy in all duplications on the evolutionary path between the two.

Our method is based on the clustering of genes based on their genomic context; two genes that have very similar gene sets in their neighborhood are more likely to be positional orthologs. Let *A* = (*a*_1_, …, *a*_*n*_) denote the gene order of a circular chromosome, where *a*_*i*_ is the identifier of the homolog group of the *i*-th gene in the chromosome. Fix a position *i* in the gene order. Consider the size *k* + 1 set of homology group identifiers to the left of and including *a*_*i*_. We refer to this set, without *a*_*i*_, as the *left k-local context* of the gene *i*. Analogously, we define the right *k*-local context. The *k-local context* of gene *i* is defined as the union of its left and right *k*-local contexts. For linear chromosomes, the left and right contexts may contain fewer than *k* elements if the position is near an extremity. In this context, we can also refer to *k* as the *radius* of the local context. We define the *k-local context similarity* between two genes as the Jaccard similarity between their *k*-local contexts.

As an example, consider the gene order (2, 1, 3, 3, 4, 5, 3, 5, 7, 2, 8, 2, 4, 10, 1). The left 3-local context of the 6th gene (with homolog group 5) is {4, 3, 1}, and its right 3-local context is {3, 7, 2}. Note that 5 is skipped in the right context, as it matches the homolog group of the gene under consideration. The combined 3-local context is therefore {1, 2, 3, 4, 7}. The 3-local context of the 8th gene (also from homolog group 5) is {1, 2, 3, 4, 7, 8}. Their 3-local context similarity is thus 5*/*6.

To break ties between gene pairs with identical local context similarity, we use a secondary similarity measure based on sequence similarity. Let *a* and *b* denote the concatenated exon sequences of the two genes. Let *BS*(*a, b*) be the bit-score of the best BLAST [6] alignment between *a* and *b*. Define the reciprocal score between *a* and *b* as

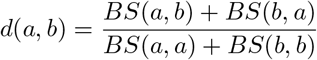

if the denominator is positive, and zero otherwise. We use the reciprocal score as the *sequence similarity* of the two genes.

The *composite similarity score* of two genes within a homolog group is the ordered pair consisting of their *k*-local context similarity and their sequence similarity. Most of the methods we apply later operate on scalar similarity scores. To this end, we define the *linearized similarity score* between two genes as 10*S* · *x* + *y*, where (*x, y*) is the composite similarity score and *S* is the sum of all sequence similarity scores between pairs of genes in that homolog group.

We formalize the problem of identifying positional orthologs as a maximization problem we call the MAXIMUM COLORFUL GRAPH PARTITION problem.

### 1.2 Maximum Colorful Graph Partition

Consider a graph *G* = (*V, E*) with a non-negative edge weight function *w*: *E* → **R**_+_ and a vertex labeling function *λ*: *V* → **N**. Let *X* ⊆ *V* be a subset of vertices, and let *G*_*X*_ be the corresponding induced subgraph (i.e., *G*_*X*_ contains all vertices from *X* and all edges of *G* that connect vertices in *X*). We denote the sum of the weights of edges in *G*_*X*_ as

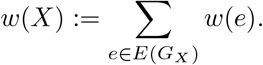

We call a subset *X colorful* if all labels of its vertices are distinct. We call a partition Ξ of the vertices *V colorful* if all its subsets are colorful. We denote the total weight of a partition Ξ as

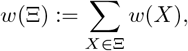

and refer to this quantity as the *weight of the partition* Ξ. The individual sets in the partition are called *clusters*.

#### Definition 1

(Maximum Colorful Graph Partition (MCGP)). *Given a graph G* = (*V, E*) *with a non-negative edge weight function w*: *E* → **R**_+_ *and a vertex labeling function λ*: *V* → **N**, *find a colorful partition* Ξ *of V that maximizes the total weight w*(Ξ).

In our setting, the vertices represent individual genes from a given homolog group, the labeling function maps each gene to its organism, and the edge weight function assigns the linearized similarity score between the corresponding genes. Each set in the resulting colorful partition then corresponds to a cluster of positional orthologs.

Note that the MCGP problem—also referred to as the Minimum Orthogonal Partition problem by He at al. [11]—is equivalent to finding a maximum-weight *clique cover* in a *k*-partite graph, where *k* is the number of distinct organisms and each partition class corresponds to a color (organism). This equivalent formulation is called the Maximum Disjoint *k*-Clique problem by [11]. Unweighted versions of this problem have been previously studied under the terms *colorful components* [5, 2] *and colorful orthology clustering* [20].

## 2 Solutions for maximum colorful graph partition problem

The MCGP problem can be expressed as an integer linear programming (ILP) maximization task. The objective function is the sum of the weights of the edges within the selected clusters. The constraints must prevent vertices with the same label from appearing in the same cluster. We present two exact ILP formulations of the MCGP problem, along with one ILP-based heuristic.

To simplify the notation, we introduce several technical assumptions. Let the vertices *V* be identified with the integers {1, …, |*V*|}. Let the set of edges *E* consist of ordered pairs (*i, j*) such that *i < j*. Define *Ē* as the set containing both (*i, j*) and (*j, i*) for every edge (*i, j*) ∈ *E*. We assume that no edge connects two vertices with the same label, i.e., for all (*i, j*) ∈ *E, λ*(*i*)≠ *λ*(*j*).

### 2.1 Exact ILP formulation using all-pairs shortest path

We can interpret the MCGP problem as the task of selecting the heaviest subset of edges such that no two vertices with the same label are in the same connected component. These connected components correspond to the resulting clusters. We use standard ILP techniques related to all-pairs shortest path.

Let the binary variable *D*_*i,j*_ indicate whether edge (*i, j*) is included in the solution. The objective function is then given by:

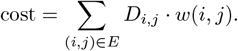

Let the binary variable *C*_*i,j*_ indicate whether vertices *i* and *j* are connected (possibly via a path) in the selected subgraph. To prevent vertices with the same label from being in the same component, we enforce:

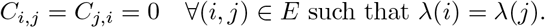

The following defines the semantics of the *C*_*i,j*_ variables using ILP constraints.

We use a construction similar to the Floyd-Warshall algorithm. Let the binary variable *C*_*i,j,k*_ indicate whether there is a path between vertices *i* and *j* using only selected edges, such that all intermediate vertices have indices at most *k*. We refer to such paths as *k-paths*. For the base case *k* = 0, the only allowed paths are single edges without intermediaries:

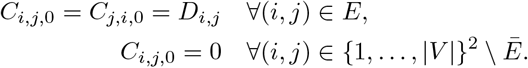

For *k >* 0, we use the recurrence:

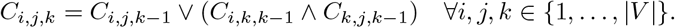

Finally, the connectivity variables are defined by:

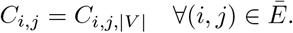

The total number of decision variables is therefore *O*(|*V*| ^3^), and the number of constraints is also *O*(|*V*| ^3^), with the total number of non-zero entries in the constraint matrix being *O*(|*V*| ^3^).

Logical AND and OR between binary variables are implemented using standard ILP encodings provided by the Gurobi API (see Appendix for details).

### 2.2 Exact ILP formulation using cluster variables

Let a discrete variable *P*_*i*_ ∈ {1, …, |*V*|} represent the cluster to which vertex *i* belongs. Let a binary variable *D*_*i,j*_ indicate whether vertices *i* and *j* belong to the same cluster. If they do, the edge between them contributes to the total weight of the partition:

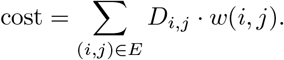

Let *Q*_*i,j*_ := |*P*_*i*_ −*P*_*j*_| be a discrete variable representing the absolute difference in cluster assignments. This absolute value is implemented using standard ILP constructs provided by the Gurobi API (see Appendix).

Then, vertices *i* and *j* are in different clusters if *Q*_*i,j*_ is strictly positive:

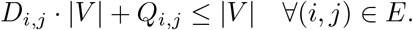

The color constraint is enforced by ensuring that no two vertices of the same label can belong to the same cluster:

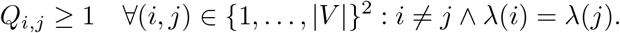

The total number of decision variables is *O*(|*V*|^2^), the number of constraints is *O*(|*V*|^2^), and the total number of non-zero elements in the constraint matrix is *O*(|*V*|^2^).

### 2.3 Heuristic ILP formulation using star cover

A star is a complete bipartite graph *K*_1,∗_, where the first part consists of exactly one vertex. We refer to that vertex as the star’s *center*. In this heuristic, our goal is to cover the input graph with such stars instead of complete cliques. The key difference is that we maximize only the sum of the edge weights *within the stars*.

We formulate the constraints as follows. Each edge can be selected into the solution *with a direction*. Let a binary variable *C*_*i,j*_ indicate that the directed edge *i* →*j* is included in the solution. We impose a constraint that an edge can be selected in at most one direction:

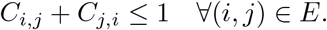

The objective function is:

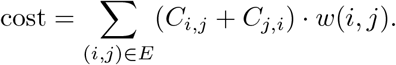

Each selected edge must *point toward the center* of its respective star. This leads to the following constraint: a vertex is either not the center of a star (and thus may have at most one outgoing edge and no incoming edges), or it is the center (with unrestricted incoming edges and no outgoing ones):

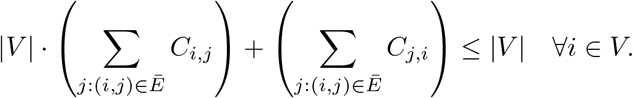

To enforce colorful clusters, we ensure that each vertex has at most one incoming edge from a given color:

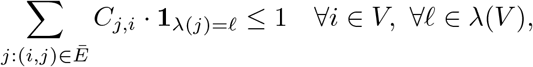

where the Iverson bracket **1**_*A*_ equals 1 if the logical condition *A* is true, and 0 otherwise. Here, |*λ*(*V*)| denotes the number of distinct labels (colors) in the input graph.

The total number of decision variables is *O*(|*E*|), the number of constraints is *O*(|*E*| + |*V* | · |*λ*(*V*)|), and the total number of non-zero entries in the constraint matrix is *O*(|*E*|).

### 2.4 Greedy Algorithm #1

In simple terms, this algorithm adds edges to the solution as long as doing so does not connect two vertices of the same color.

Sort the edges by their weight in descending order. Then, one by one, check whether adding a given edge would connect two vertices of the same color. If not, include the edge in the solution.

### 2.5 Greedy Algorithm #2: universal polishing

This iterative heuristic improves a given solution by locally reassigning vertices of a chosen color using maximum weighted matching.

In each iteration, the algorithm picks a random color, disconnects all vertices of that color from their current clusters, and finds the optimal way to reconnect them.

To ensure termination, we maintain a list of unprocessed colors. At each step, a new color is selected from this list. If reassigning the selected color results in a better solution, all colors are restored to the list for potential future reassignment. If not, the color is removed from the list. The algorithm terminates when the list of unprocessed colors is empty, returning the best solution found.

This algorithm is guaranteed to produce a solution with cost at least as good as the input. It is especially effective as a lightweight polishing step following more aggressive or approximate algorithms.

### 2.6 He et al. (2004) algorithm with guide tree

He et al. [11] proposed an approximation algorithm for MCGP. In each iteration, two labels (colors) are selected, and vertices of those labels are merged into clusters using maximum weighted matching. The resulting clusters (of size 1 or 2) are then contracted into single vertices with a new common label. Edge weights between the new vertices and the rest of the graph are updated by summing the weights of the corresponding original edges. This reduces the number of distinct colors by one in each iteration.

Zheng et al. [26] suggested using a guide tree, where pairs of closest colors (according to a phylogeny) are merged first. We adopt this variant of the algorithm.

### 2.7 Connectivity pruning

We determine a weight threshold such that removing all edges with weights below this threshold does not disconnect the input graph. This preprocessing step helps reduce the size of the ILP instance by pruning low-weight, redundant edges.

## 3 Experiments

### 3.1 Large gene families from the *Anopheles* genus

We selected four genomes from the *Anopheles* genus: *A. albimanus, A. atroparvus, A. merus*, and *A. stephensii*. From these, we randomly chose several small gene families and also included the 20 largest gene families (by number of genes). We excluded the ILP algorithm based on the Floyd–Warshall construction due to its excessive memory consumption.

All experiments were run on a local machine with an Intel(R) Xeon(R) W-2245 CPU @ 3.90GHz, using 8 threads and a time limit of 86 400 seconds (i.e. one day).

Table 1 shows that all algorithms achieve similar partition weights. It also demonstrates that using Greedy Algorithm #2 as a post-processing polishing step has a negligible impact on solution quality. Additionally, the pruning strategy does not significantly degrade solution quality, while consistently reducing the running time (though the effect is not always pronounced), as seen in Table 2.

**Table 1.** Relative efficiency of our algorithms for the MCGP problem on gene families from four *Anopheles* genomes. Relative efficiency is computed as the fraction of the best-known score for each task, including solutions refined by Greedy Algorithm #2.

| Task | Nodes | ILP-components | ILP-components-pruned | greedy #1 | greedy #2 | He2004 | ILP-star |
| --- | --- | --- | --- | --- | --- | --- | --- |
| anopheles-016-1082 | 16 | 1.00 | 1.00 | 1.00 | 1.00 | 1.00 | 1.00 |
| anopheles-017-1268 | 17 | 1.00 | 1.00 | 1.00 | 1.00 | 1.00 | 1.00 |
| anopheles-018-6369 | 18 | 1.00 | 1.00 | 0.99 | 0.97 | 1.00 | 0.90 |
| anopheles-021-1548 | 21 | 1.00 | 1.00 | 0.99 | 1.00 | 1.00 | 0.98 |
| anopheles-022-4178 | 22 | 1.00 | 1.00 | 1.00 | 1.00 | 1.00 | 0.98 |
| anopheles-024-912 | 24 | 1.00 | 1.00 | 1.00 | 1.00 | 1.00 | 0.95 |
| anopheles-028-3707 | 28 | 1.00 | 1.00 | 1.00 | 1.00 | 1.00 | 1.00 |
| anopheles-032-6396 | 32 | 1.00 | 1.00 | 0.98 | 1.00 | 0.99 | 0.96 |
| anopheles-035-1019 | 35 | 1.00 | 1.00 | 0.92 | 1.00 | 1.00 | 0.97 |
| anopheles-036-2530 | 36 | 1.00 | 1.00 | 1.00 | 1.00 | 1.00 | 0.99 |
| anopheles-036-3609 | 36 | 1.00 | 1.00 | 0.99 | 1.00 | 0.97 | 0.97 |
| anopheles-036-5419 | 36 | 1.00 | 0.99 | 0.99 | 1.00 | 1.00 | 0.96 |
| anopheles-041-8053 | 41 | 1.00 | 1.00 | 1.00 | 1.00 | 1.00 | 1.00 |
| anopheles-043-411 | 43 | 0.00 | 0.99 | 1.00 | 0.96 | 0.96 | 0.96 |
| anopheles-043-988 | 43 | 1.00 | 1.00 | 0.99 | 1.00 | 0.99 | 0.97 |
| anopheles-047-3095 | 47 | 1.00 | 1.00 | 0.99 | 1.00 | 0.99 | 0.99 |
| anopheles-050-3212 | 50 | 1.00 | 1.00 | 0.91 | 1.00 | 0.94 | 0.91 |
| anopheles-051-13657 | 51 | 1.00 | 1.00 | 1.00 | 1.00 | 1.00 | 1.00 |
| anopheles-053-4999 | 53 | 1.00 | 1.00 | 1.00 | 1.00 | 0.91 | 1.00 |
| anopheles-055-850 | 55 | 1.00 | 1.00 | 0.94 | 1.00 | 0.99 | 0.94 |
| anopheles-059-1281 | 59 | 1.00 | 1.00 | 0.99 | 1.00 | 1.00 | 0.99 |
| anopheles-060-1912 | 60 | 1.00 | 1.00 | 0.97 | 1.00 | 0.98 | 0.94 |
| anopheles-062-932 | 62 | 1.00 | 1.00 | 1.00 | 0.97 | 1.00 | 0.95 |
| anopheles-068-1499 | 68 | 1.00 | 1.00 | 0.96 | 0.99 | 0.97 | 0.93 |
| anopheles-074-189 | 74 | 0.99 | 0.00 | 0.86 | 0.99 | 0.98 | 0.90 |
| anopheles-136-404 | 136 | 0.98 | 0.99 | 0.96 | 1.00 | 0.99 | 0.96 |
| anopheles-206-133 | 206 | 0.99 | 0.98 | 0.97 | 0.99 | 0.99 | 0.98 |

**Table 2.** Running time (in seconds) of our algorithms for the MCGP problem on gene families from four *Anopheles* genomes. Note that the time limit was set to 86 400 seconds (1 day).

| Task | Nodes | ILP-components | ILP-components-pruned | greedy #1 | greedy #2 | He2004 | ILP-star |
| --- | --- | --- | --- | --- | --- | --- | --- |
| anopheles-016-1082 | 16 | 0 | 0 | 0 | 0 | 1 | 0 |
| anopheles-017-1268 | 17 | 0 | 0 | 0 | 0 | 1 | 0 |
| anopheles-018-6369 | 18 | 3 | 3 | 0 | 0 | 0 | 0 |
| anopheles-021-1548 | 21 | 2 | 2 | 0 | 0 | 0 | 0 |
| anopheles-022-4178 | 22 | 1 | 1 | 0 | 0 | 0 | 0 |
| anopheles-024-912 | 24 | 1 | 0 | 0 | 0 | 1 | 0 |
| anopheles-028-3707 | 28 | 0 | 1 | 0 | 0 | 1 | 0 |
| anopheles-032-6396 | 32 | 34222 | 24651 | 0 | 1 | 1 | 0 |
| anopheles-035-1019 | 35 | 248 | 155 | 0 | 1 | 1 | 1 |
| anopheles-036-2530 | 36 | 5 | 3 | 0 | 0 | 1 | 0 |
| anopheles-036-3609 | 36 | 130 | 56 | 0 | 0 | 1 | 0 |
| anopheles-036-5419 | 36 | 869 | 2 | 0 | 0 | 1 | 0 |
| anopheles-041-8053 | 41 | 35 | 0 | 0 | 1 | 1 | 0 |
| anopheles-043-411 | 43 | 367 | 165 | 0 | 0 | 1 | 1 |
| anopheles-043-988 | 43 | 699 | 574 | 0 | 1 | 1 | 0 |
| anopheles-047-3095 | 47 | 86433 | 86435 | 0 | 1 | 1 | 0 |
| anopheles-050-3212 | 50 | 60629 | 1641 | 0 | 1 | 1 | 0 |
| anopheles-051-13657 | 51 | 13 | 4 | 0 | 0 | 1 | 0 |
| anopheles-053-4999 | 53 | 39 | 2 | 0 | 1 | 0 | 0 |
| anopheles-055-850 | 55 | 86408 | 86408 | 0 | 1 | 1 | 1 |
| anopheles-059-1281 | 59 | 86402 | 86402 | 0 | 1 | 1 | 0 |
| anopheles-060-1912 | 60 | 86408 | 86402 | 0 | 1 | 1 | 0 |
| anopheles-062-932 | 62 | 23 | 9 | 0 | 1 | 1 | 0 |
| anopheles-068-1499 | 68 | 86409 | 86403 | 0 | 2 | 1 | 0 |
| anopheles-074-189 | 74 | 86407 | 86403 | 0 | 1 | 1 | 1 |
| anopheles-136-404 | 136 | 86402 | 86401 | 0 | 5 | 1 | 2 |
| anopheles-206-133 | 206 | 86404 | 86411 | 0 | 9 | 2 | 1 |

Table 2 also illustrates that the greedy strategies achieve comparable solution quality while being drastically faster than the exact ILP-based algorithms.

Figure 1, which depicts the best solution found over time for the largest gene family anopheles-206-133 (with 206 nodes), shows that it takes several thousand seconds (i.e., hours), even with multiple threads, for the exact ILP-based algorithm to reach a solution within 95% of the cost of a greedy solution.

**Figure 1.**
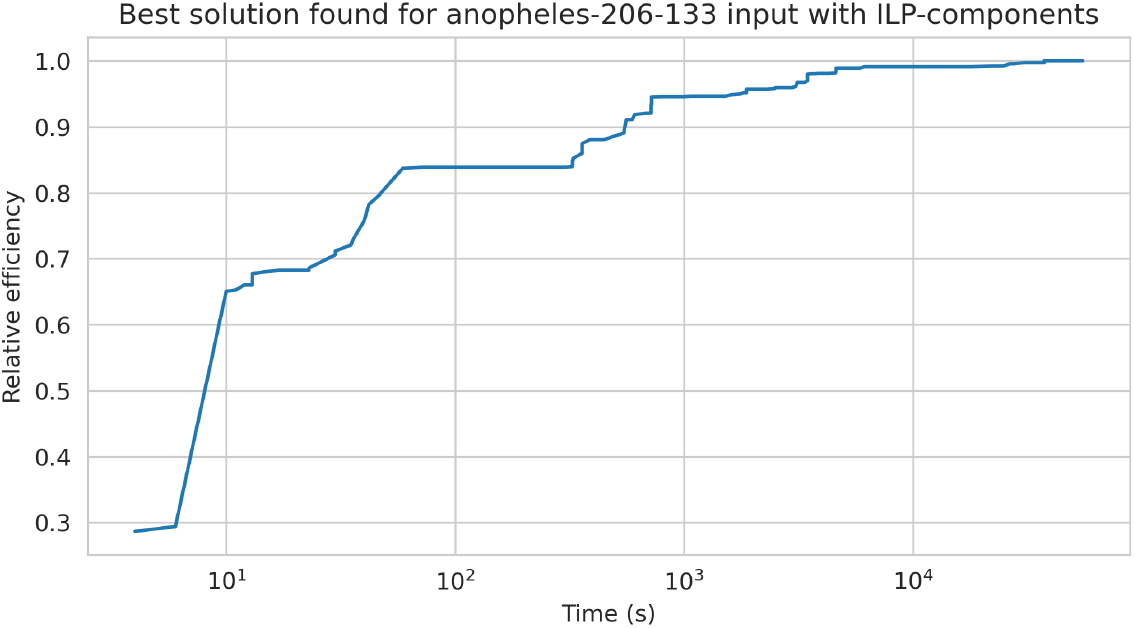
Best solution found over time by the ILP-components algorithm on the `anopheles-206-133` gene family (206 nodes). The horizontal axis is in logarithmic scale.

### 3.2 Synthetic genomes using Zombi

We simulated the evolution of 10 species with 100 gene families using the `Zombi` simulation tool [7]. `Zombi` is a comprehensive simulator that generates a species tree, simulates gene order evolution along the tree, and finally produces nucleotide sequences of the resulting genes trees. We used the default parameters of `Zombi` in most cases, varying only the duplication rate (called `dup_rate`) from 1.0 and 8.0. Duplications and losses (rate of 1) modified the gene content while inversions (rate of 2) and duplications modified the gene neighborhoods. The nucleotide length of individual genes was set to 1 500 bp, and the `K2P` model was used for simulating gene sequence evolution.

To measure the similarity between the true gene family partition and the results produced by our algorithms, we used the Rand index. It is computed as follows. Let *A* and *B* be two partitions of a set *X*. Let *x* be the number of element pairs in *X* that appear in the same cluster in both *A* and *B*, and let *y* be the number of pairs that appear in different clusters in both *A* and *B*. Then, the Rand index of *A* and *B* is defined as

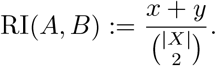

To assess the inherent difficulty of a gene family partitioning task in terms of the Rand index, we devised a simple *random* baseline strategy. Given a gene family with *k* resulting clusters, we create *k* buckets and randomly distribute the genes from each organism among them, placing at most one gene per organism in each bucket. Note that the random partition has an advantage, in that it assumes knowledge of the true number of clusters! This is a distinct advantage for the random clustering when recent gene loss has occurred in duplicated gene families.

Our ILPs were run with a time limit of 2 minutes per gene family, while the `FFGC` time limit was set to 5 minutes per pair of genomes (rather than the default 120 used for the *Anopheles* data).

Our experiments show that the heuristic star cover ILP formulation performs worse than both the algorithm of He et al. [11] and the exact ILP formulation using cluster variables (see Figure 2). We therefore exclude the star-cover-based heuristic from further analysis.

**Figure 2.**
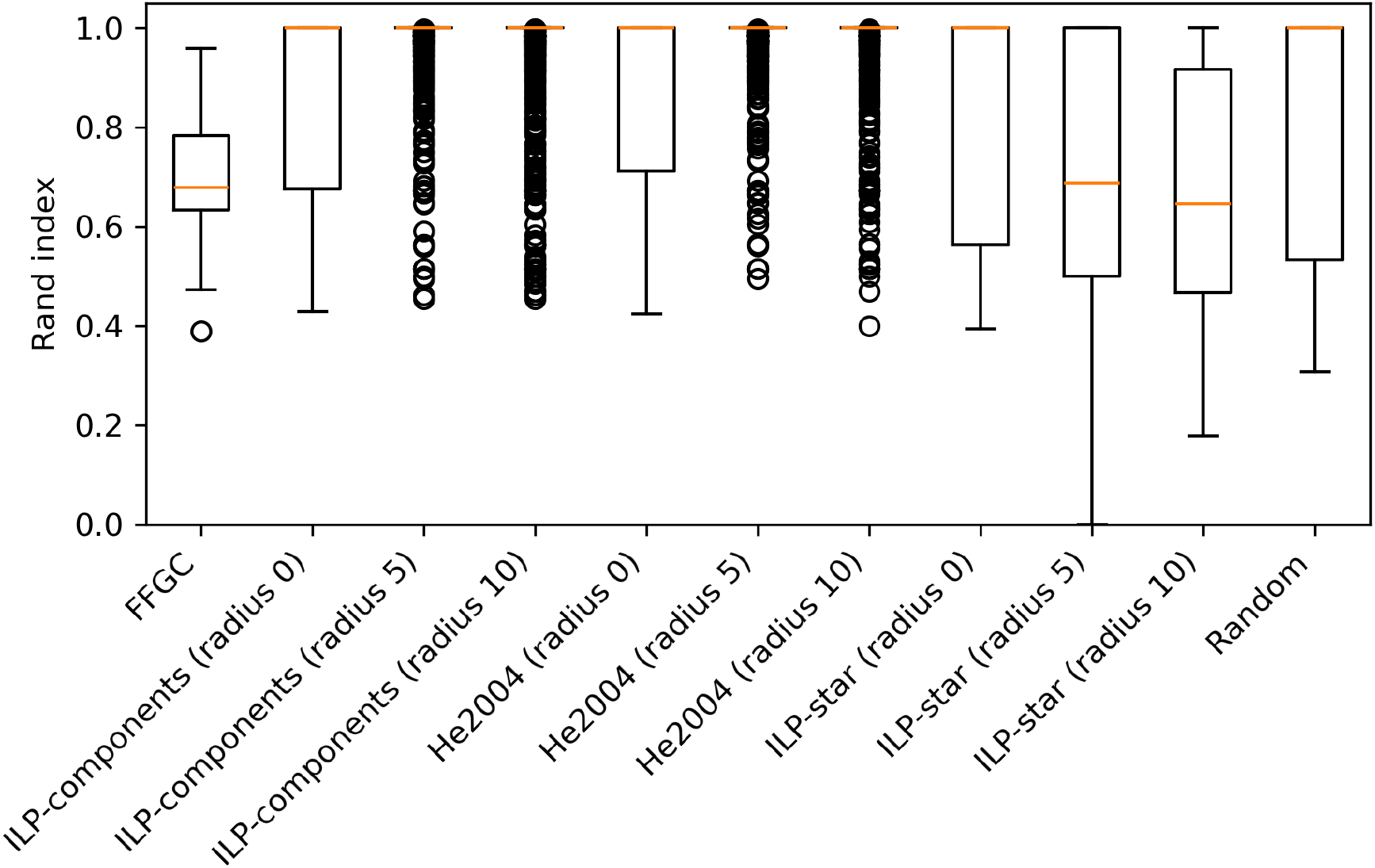
Genomic context matters. Boxplots of Rand index values over three neighborhood radius values are shown for all duplication rates combined. Use of the neighboorhood is essential to improved performance.

Figure 2 shows the importance of using the gene neighborhood, as algorithms perform worse when the local context radius is set to zero.

The experiments also show that the `FFGC` algorithm under-performs. This behavior persists even when substitution mutations in the gene sequences are eliminated (see Figure 5 in the Supplement). To further investigate this issue, we computed the median Rand index for gene families grouped by different gene copy numbers (see Figure 3). The results indicate that `FFGC` performs particularly poorly on gene families with low copy numbers, by essentially assigning genes as insertions/deletions rather than keeping them in a family. This alarming fact lead us to investigate this deficiency, and we found that BLAST similarity scores were not being computed for dissimilar sequences. We lowered the nucleotide substitution rate to 0 and found a similar pattern (see Figure 5). It remains unclear as to whether this performance is due to an implementation issue, or rather a deficiency in the method itself.

**Figure 3.**
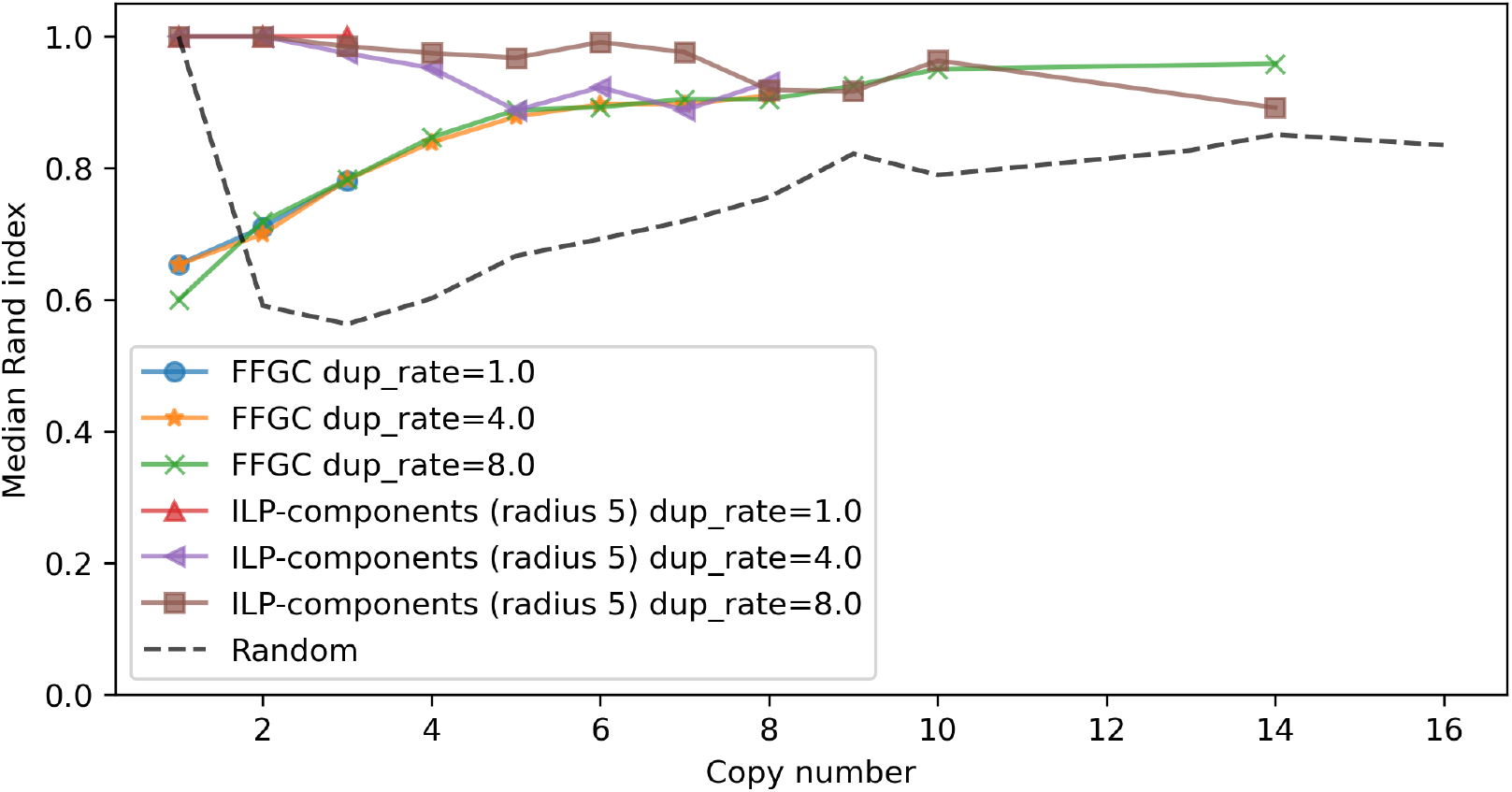
Median Rand index of `FFGC` for varying gene copy numbers. Lines terminate since low duplication rates yield low copy numbers. The dashed line represents the random across all simulator settings, noting that the random method knows the true number of clusters.

The results also show that the algorithm of He et al.[11] and our exact ILP solution perform similarly well across gene families with varying copy numbers, except when deprived of local context information (see Figure 4).

**Figure 4.**
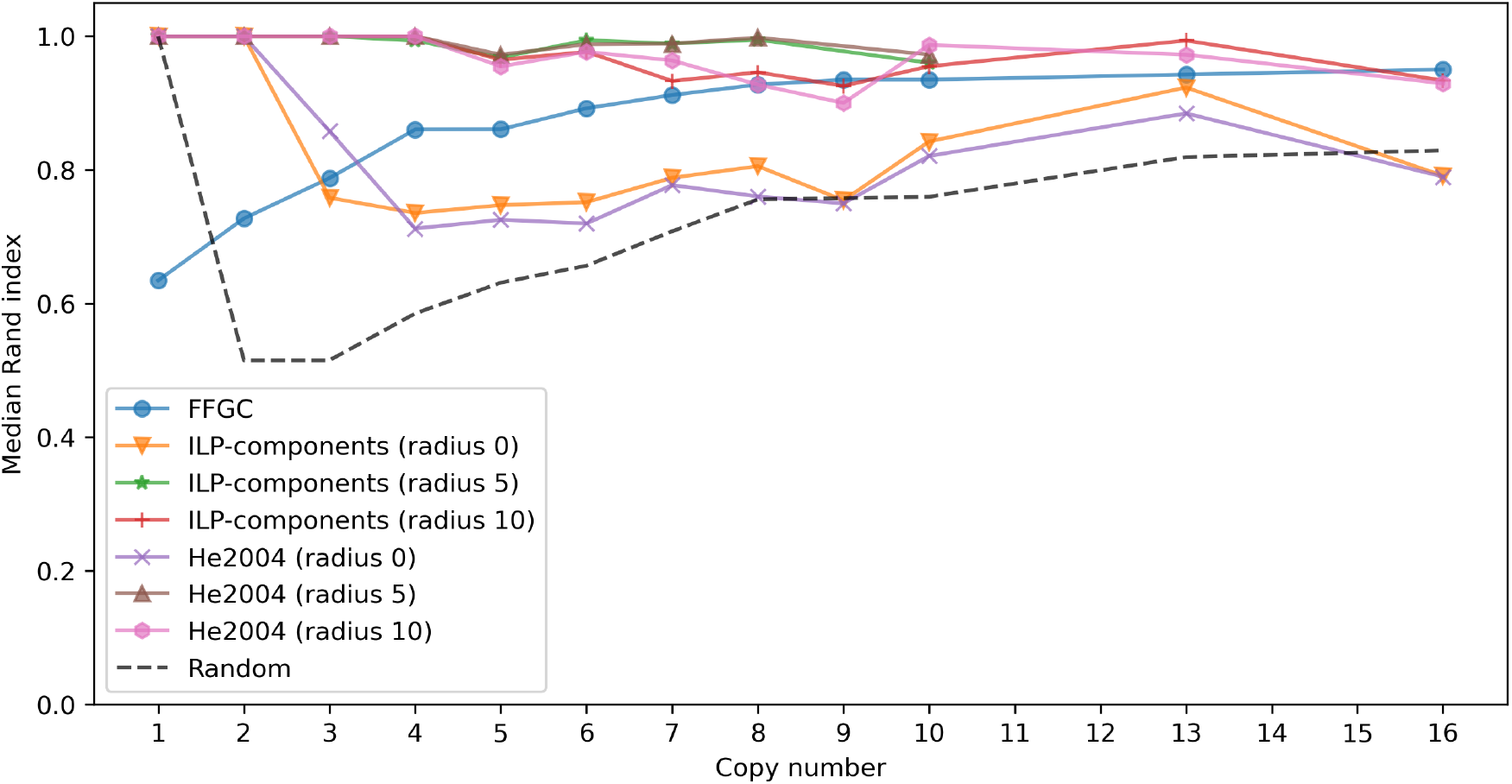
Median Rand index by gene copy number, for `dup_rate = 6.` Gene neighborhood radius improves inference.

**Figure 5.**
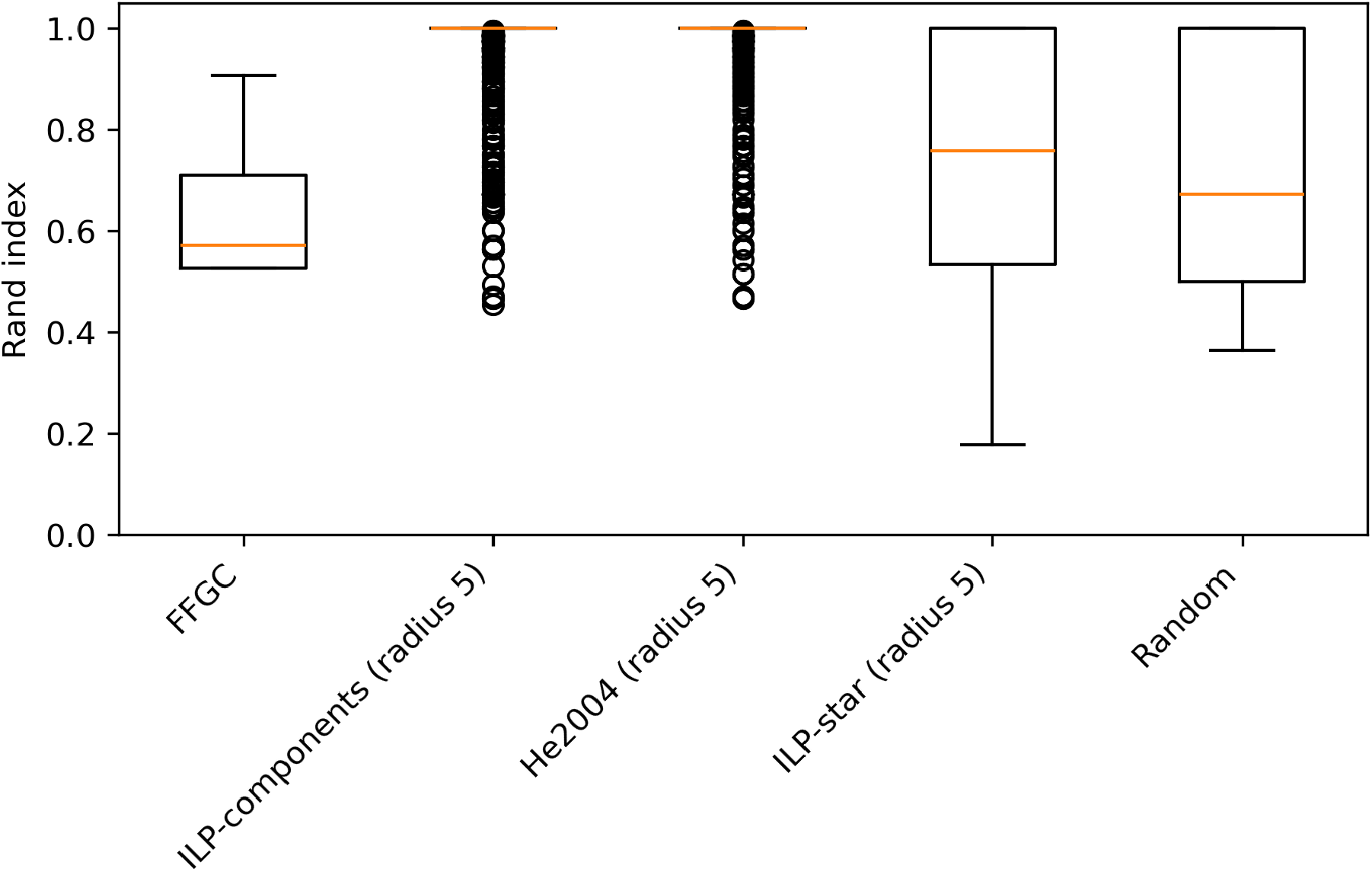
Boxplots of Rand index values for individual gene families in synthetic experiments with different duplication rates *and zero substitution rate*.

Moreover, increasing the local context radius does not lead to a notable improvement in performance. This may be explained by the relatively small size of the simulated genome (only 100 genes), where larger local context windows do not contribute additional useful information.

## 4 Conclusion

We introduced a novel mathematical model for the identification of positional orthologs, and proposed several exact and heuristic algorithms, including an adaptation of the approximation algorithm of He et al. [11]. We evaluated the performance of our methods on large gene families from the *Anopheles* genus.

Additionally, we compared the ability of our algorithms to identify positional orthologs against the established `FFGC` tool using synthetic datasets with known ground truth. The results demonstrate that our model can reliably identify positional orthologs, with two algorithms—namely, the exact ILP formulation using cluster variables and the He et al. [11] heuristic—emerging as the most effective.

Since our mathematical model treats each gene family independently, a hybrid approach combining multiple algorithms may be employed for improved performance. Moreover, our approach appears to outperform the existing `FFGC` tool in terms of both accuracy and robustness. Note that we *do not* draw the conclusion that `FFGC` cannot be suited to this task, as it is unclear if the current performance is a consequence of implementation, rather than methodology.

Most crucially, by comparing MCGP solutions with and without genomic context on simulated data, we demonstrated that the use of context is essential for the identification of positional orthologs.

## Availability

Our software is available at https://github.com/japdlsd/positional_homologs.

## A Standard ILP constructions

### A.0.0.1 AND construction

Let *X, Y*, and *Z* be binary variables with the following constraints:

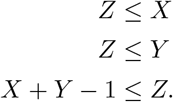

Then, *Z* = *X* ∧ *Y*.

### A0.0.2 OR construction

Let *X, Y*, and *Z* be binary variables with the following constraints:

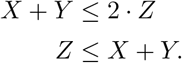

Then, *Z* = *X* ∨ *Y*.

### A0.0.3 Absolute difference construction

Let *X, Y* be discrete variables with range {1, …, *m*}, *W* be an auxiliary binary variable, and *Z* be a discrete variable with the following constraints:

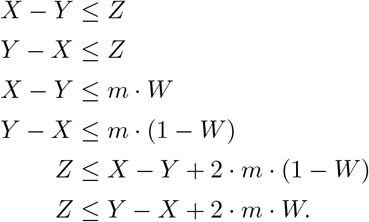

Then, *Z* = |*X* −*Y*|. Also, *W* has value 1 if *X > Y* and value 0 if *X < Y*, and whichever value in case *X* = *Y*.

## B Additional figures

